# Dosage of duplicated and antifunctionalized homeobox proteins influences spikelet development in barley

**DOI:** 10.1101/2021.11.08.467769

**Authors:** Venkatasubbu Thirulogachandar, Geetha Govind, Götz Hensel, Sandip M. Kale, Markus Kuhlmann, Lennart Eschen-Lippold, Twan Rutten, Ravi Koppolu, Jeyaraman Rajaraman, Sudhakar Reddy Palakolanu, Christiane Seiler, Shun Sakuma, Murukarthick Jayakodi, Justin Lee, Jochen Kumlehn, Takao Komatsuda, Thorsten Schnurbusch, Nese Sreenivasulu

## Abstract

Illuminating the mechanisms of inflorescence architecture of grain crops that feed our world may strengthen the goal towards sustainable agriculture. Lateral spikelet development of barley (*Hordeum vulgare* L.) is such an example of a floral architectural trait regulated by VRS1 (Vulgare Row-type Spike 1 or Six-rowed Spike 1, syn. HvHOX1). Its lateral spikelet-specific expression and the quantitative nature of suppressing spikelet development were previously shown in barley. However, the mechanistic function of this gene and its paralog HvHOX2 on spikelet development is still fragmentary.Here, we show that these duplicated transcription factors (TFs) have contrasting nucleotide diversity in various barley genotypes and several Hordeum species. Despite this difference, both proteins retain their basic properties of the homeodomain leucine zipper class I family of TFs. During spikelet development, these genes exhibit similar spatiotemporal expression patterns yet with anticyclic expression levels. A gene co-expression network analysis suggested that both have an ancestral relationship but their functions appear antagonistic to each other, i.e., HvHOX1 suppresses whereas HvHOX2 rather promotes spikelet development. Our transgenic promoter-swap analysis showed that HvHOX2 can restore suppressed lateral spikelets when expression levels are increased; however, at its low endogenous expression level, HvHOX2 appears dispensable for spikelet development. Collectively, this study proposes that the dosage of the two antagonistic TFs, HvHOX1 and HvHOX2, influence spikelet development in barley.

## Introduction

Cereals such as maize (*Zea mays* L.), rice (*Oryza sativa* L.), wheat (*Triticum* spp.), and barley (*Hordeum vulgare* L.) are major grass species that feed most of the population on earth. Understanding the genetic regulation of inflorescence (flower-bearing structure) architecture in these cereal crops may shed light on the basic developmental patterning of floral meristems and reveal potential pathways to improve their yield. Barley, along with other major cereal crops (wheat, rye, and triticale) belonging to the Triticeae tribe, possesses a branchless inflorescence known as ‘spike’ (Ullrich, 2011; Koppolu and Schnurbusch, 2019). In general, a barley spike forms three spikelets on its rachis (inflorescence axis) nodes – one central and two lateral spikelets in an alternating, opposite arrangement (distichous) (Bonnett, 1935; Koppolu and Schnurbusch, 2019; Zwirek et al., 2019). The spikelet, a small/condensed spike, is considered the basic unit of the grass inflorescence (Clifford et al., 1987; Kellogg et al., 2013). A barley spikelet forms a single floret that is subtended by a pair of glumes. Typically, a barley floret consists of one lemma, one palea, two lodicules, three stamens, and a monocarpellary pistil (i.e., single carpel) (Waddington et al., 1983; Forster et al., 2007). Based on the fertility of the lateral spikelets/florets, barley is classified into two- and six-rowed spike types. In two-rowed types, the lateral spikelets are smaller (compared to the central spikelets), awnless (extension of the lemma is absent), and sterile, while the central spikelets are bigger, awned, and fertile. In six-rowed types, both the lateral and central spikelets are awned and fertile.

The major gene responsible for the lateral spikelet fertility was found to be a homeodomain leucine zipper class I (HD-ZIP I) transcription factor, known as *VRS1* (***V****ulgare* ***R****ow-type* ***S****pike1* or *Six-rowed Spike 1*, syn *HvHOX1*) (Komatsuda et al., 2007). Transcripts and proteins of *HvHOX1* had previously been found in barley spikes, predominantly in the lateral florets and most strongly in the carpels, corroborating a role of HvHOX1 as negative regulator of lateral floret development and fertility (Komatsuda et al., 2007; Sakuma et al., 2010; Sakuma et al., 2013). Recently, a very similar function has also been identified for its orthologous wheat gene during apical floret abortion(Sakuma et al., 2019). In recent years, *HvHOX1* was shown to be also expressed in other organs, such as leaves, where in analogy to its effects on lateral spikelet development, it negatively affects the size of leaf primordia and results in narrower leaves in two-rowed barleys (Thirulogachandar et al., 2017). Further supporting its suppressive function, one specific allele of *HvHOX1* is responsible for the extremely reduced lateral spikelet size in *deficiens* barley (Sakuma et al., 2017). Interestingly, *HvHOX2*, the paralog of *HvHOX1*, was also identified in barley. Although *HvHOX2* is expressed in a wide variety of organs including leaves, coleoptile, root, and spike; tissue-wise, it is mainly found in vascular regions particularly those at the base of lateral spikelets (pedicel) and rachis, thus suggesting a role in the promotion of development (Sakuma et al., 2010; Sakuma et al., 2011; Sakuma et al., 2013). In addition to *HvHOX1*, four other genes, *VRS2, VRS3, VRS4*, and *VRS5* or *INT-C* (*intermedium-spike c*), were reported to be involved in the suppression of lateral spikelet fertility (Ramsay et al., 2011; Koppolu et al., 2013; Bull et al., 2017; van Esse et al., 2017; Youssef et al., 2017). Notably, VRS4, the ortholog of maize RAMOSA2 (RA2) appeared to be functionally upstream of *HvHOX1* but not of *HvHOX2* (Koppolu et al., 2013; Sakuma et al., 2013). Later, VRS3 was also identified as an upstream regulator of *HvHOX1*, and in certain stages also of *HvHOX2* (Bull et al., 2017; van Esse et al., 2017).

Despite the detailed studies on *HvHOX1*’s expression pattern and mutants, the mechanistic role of HvHOX1 on barley spikelet development is still unclear. The same holds true for HvHOX2 while its suggested role in barley development has yet to be validated(Sakuma et al., 2010; Sakuma et al., 2013). In this study, we show that HvHOX1 and HvHOX2 proteins are functional HD-ZIP class I transcription factors. Our transcript expression studies suggest that both have similar spatiotemporal expression patterns; however, with a contrasting dosage of transcripts in central and lateral spikelets during spikelet development. Based on our combined results, we conclude that both genes are ancestrally related but act antagonistically to each other, i.e., HvHOX1 suppresses whereas HvHOX2 rather promotes spikelet development. Our transgenic promoter-swap analysis shows that *HvHOX2* can restore suppressed lateral spikelets when transcript levels are increased, most likely, by modulating the adverse effects caused by *HvHOX1*. At low endogenous transcript levels, however, *HvHOX2* appears dispensable for spikelet development. Collectively, our findings recommend that *HvHOX1* and *HvHOX2* act antagonistic to each other, and that the dosage of their transcripts influences barley spikelet development.

## Results

### *HvHOX2* nucleotide diversity is highly conserved compared to its paralog *HvHOX1*

The eight different natural alleles for *HvHOX1* known so far are grouped into two-rowed (*Vrs1*.*b2, Vrs1*.*b3, Vrs1*.*b5*, & *Vrs1*.*t1*) and six-rowed alleles (*vrs1*.*a1, vrs1*.*a2, vrs1*.*a3*, & *vrs1*.*a4)* (Komatsuda et al., 2007; Sakuma et al., 2017; Casas et al., 2018). In contrast, the nucleotide diversity of *HvHOX2* is largely unknown. To fill this gap, we sequenced the *HvHOX2* promoter (one kb) and gene (including 5’ and 3’ untranslated region) in 83 diverse spring barleys (44 two-rowed and 39 six-rowed). Surprisingly, we found only four single nucleotide polymorphisms (SNPs), restricted to the promoter (two SNPs), 5’UTR (one SNP), and intron-2 (one SNP). At the same time, the coding sequence (CDS) was identical and highly conserved in all these accessions (Supplementary Table 1). We further expanded our nucleotide diversity study by sequencing the *HvHOX1* and *HvHOX2* in 24 *Hordeum* spp. (Supplementary Table 2), which showed that the non-synonymous (*Ka*) and synonymous (*Ks*) substitution values of *HvHOX1* (*Ka* = 0.028, *Ks* = 0.049) and *HvHOX2* (*Ka* = 0.008, *Ks* = 0.051). The higher *Ka* value of *HvHOX1* than that of *HvHOX2* indicates that the evolutionary speed of *HvHOX1* is much faster than that of *HvHOX2*, otherwise, *HvHOX2* has been well conserved among the *Hordeum* species (Supplementary Table 3). A subsequent comparison of the nucleotide diversity (π) of these two genes (*HvHOX2*, Chr.2H: 139932435-139953386; *HvHOX1*, Chr.2H: 581356498-581377358) in 200 domesticated barleys(Jayakodi et al., 2020) confirmed the lower nucleotide diversity (π) of *HvHOX2* compared to *HvHOX1* (Supplementary Fig. 1A). The study also revealed two major haplotypes for the *HvHOX2* genic region, whereas *HvHOX1* possesses multiple haplotypes that span the whole region analyzed (Supplementary Fig. 1B). This difference in diversity might be due to their physical location, wherein *HvHOX1* is located in the distal end of the high recombining region of chromosome 2H, while *HvHOX2* is closer to the centromeric region on 2H. Concertedly, all the above results indicate that *HvHOX2* is highly conserved compared to its paralog *HvHOX1*.

### HvHOX1 and HvHOX2 are functional HD-ZIP class I transcription factors

In general, members of the HD-ZIP family (class I to IV) of transcription factors possess a homeodomain (HD) followed by a leucine zipper motif (LZ). The LZ motif enables the dimerization of HD-ZIP proteins, which bind to their specific DNA target (cis-element) via the HD motif (Ariel et al., 2007). The HD-ZIP class I proteins - HvHOX1 and HvHOX2 show a very high sequence identity between their HD (89.3 %) and LZ (90 %) motifs. However, they have several amino acid changes across their protein with yet unknown consequences (Supplementary Fig. 2). In particular, HvHOX1 lacks a putative AHA-like motif in its C-terminus, which was predicted to be an interaction motif with the basal transcriptional machinery (Arce et al., 2011; Capella et al., 2014) (Supplementary Fig. 2). All these similarities and discrepancies paved the way to compare the functionality of these two proteins.

We assessed the dimerization properties of HvHOX1 and HvHOX2 with the bimolecular fluorescence complementation assay. *HvHOX1* and *HvHOX2* were cloned into split-Yellow Fluorescence Protein (YFP) vectors creating N-terminal c-myc-nYFP and HA-cYFP fusions. The resulting plasmids were co-transformed with a Cyan Fluorescent Protein (CFP) construct into Arabidopsis mesophyll protoplasts. The CFP served as a transformation control, accumulating in the nucleus and cytoplasm. The detection of yellow fluorescence in all four combinations indicated that the HvHOX1 and HvHOX2 proteins are able to form homo- or heterodimers (Fig. 1A). The superimposed YFP channel (dimerization) on the CFP channel (strong nuclear signal) indicated that homo- or heterodimers of both proteins are localized in the nucleus (Fig. 1A), which is in agreement with the nuclear localization signals predicted for both proteins (Sakuma et al., 2013). This localization also implied that these dimers might bind to their *cis*-elements to transactivate their downstream genes (Fig. 1A). A western blot analysis using antibodies directed against HA and c-myc epitopes confirmed that the proteins were expressed in full-length and at similar levels (Supplementary Fig. 3). Following, we verified the DNA binding properties of HvHOX1, and HvHOX2 with an electromobility shift assay (EMSA) using the *in vitro* translated proteins and experimentally verified *HD-Zip* I cis-element from Sessa et al. (1993)(Sessa et al., 1993). A clear shift of protein-DNA bands (marked with *) was detected for both proteins, especially in higher concentrations of proteins, which indicated binding to the *HD-Zip* I cis-element (Fig. 1B). The result further suggested that HvHOX1 might have a more potent DNA binding property than HvHOX2 (Fig. 1B). We then conducted a *cis*-element competition assay to evaluate the binding specificity of the proteins to the *HD-Zip* I cis-element. Intriguingly, we observed binding of HvHOX1 to *HvHOX2* promoter and mild interactions of HvHOX2 with the *HvHOX1* and *HvHOX2* promoters (Supplementary Fig. 4). This suggests that *in vivo*, HvHOX1 potentially influences *HvHOX2* expression, similarly, HvHOX2 modulates *HvHOX1* expression.

**Figure 1:**
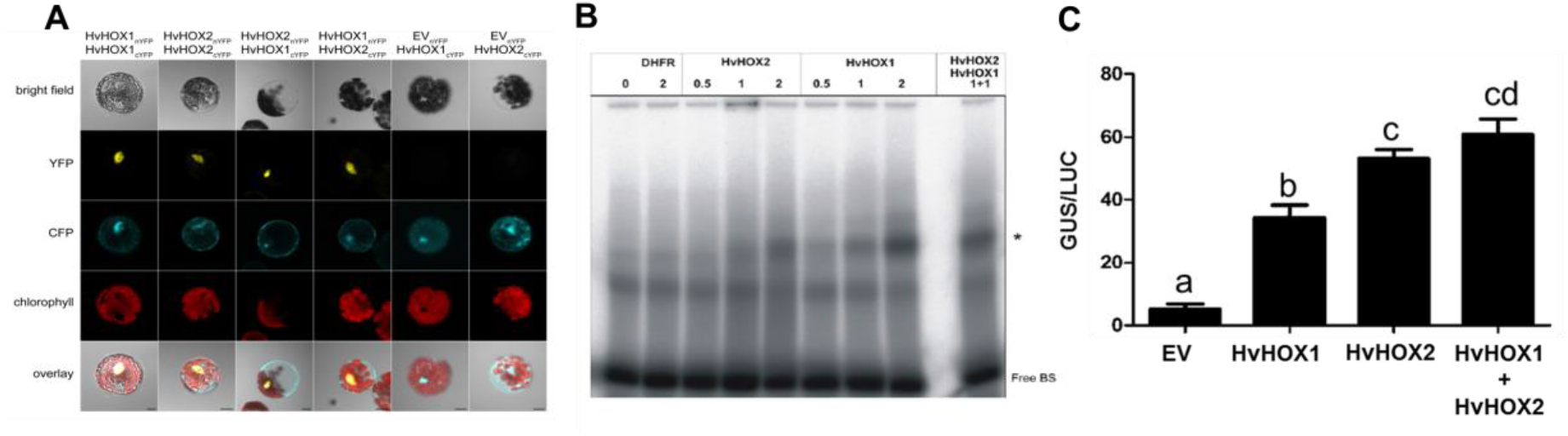
HvHOX1 and HvHOX2 are functional HD-ZIP class I transcription factors. Bimolecular fluorescence complementation assay (BiFC) for HvHOX1 and HvHOX2 proteins is shown (A). The bright field panel displays the protoplast in which the results were captured; YFP (Yellow Fluorescent Protein) panel reveals the dimer formation with the yellow color fluorescence, and CFP (Cyan Fluorescent Protein) panel discloses the location of the nucleus (blue, dark spot), and the autofluorescence of chlorophyll (red signal) is seen in the chlorophyll panel. The last overlay panel exhibits the merged signals from the above three panels. nYFP-YFP fused to N-terminal; cYFP-YFP fused to C-terminal end. Scale bar 10 µm. B) The DNA binding specificity of HvHOX1 and HvHOX2 proteins on HD-Zip I cis-element assessed by Electro Mobility Shift Assay (EMSA) is shown. Three different concentrations (0.5 µL, 1 µL, and 2 µL) of protein were used along with the DNA fragment containing the HD-Zip I cis-element (Binding sequence, BS). The shift of protein-DNA complex (*) denotes the specific DNA binding of these proteins. Also, a combination of HvHOX1 and HvHOX2 proteins (1 µL from each) also shows the protein-DNA complex. Dihydrofolate reductase (DHFR) was used as a negative control. BS-binding sequence (HD-Zip I cis element); Free BS-unbound BS; different numbers show the *in vitro* translated protein volume in µL. C) The transactivation property of HvHOX1 and HvHOX2 proteins is shown. Bar plot indicates the detected GUS activity relative to luciferase (LUC). Data shown are mean ± SE (n=3); different letters (a, b, c, and d) indicate that the mean values are significantly different at the 1% probability level, by One-way ANOVA with Newman-Keuls Multiple Comparison Test; EV-empty vector, pGAL4-4xUAS::GUS; HvHOX1-construct of GAL4-DNA binding domain fused to N-terminus of HvHOX1; HvHOX2-GAL4-DNA binding domain fused to N-terminus of HvHOX2; LUC-luciferase used for normalization; GUS-β-glucuronidase

After the dimerization and DNA binding studies, we investigated the transactivation property of these proteins *in vivo* using an Arabidopsis mesophyll protoplast system. We found that both proteins have transactivating properties, which were quantified and compared with the empty vector. Interestingly, the transactivation property of HvHOX2 was significantly higher compared to that of HvHOX1 (Fig. 1C). Collectively, all of the above results exemplified that both HvHOX1 and HvHOX2 possess DNA binding activity, can form homo- and heterodimers, and have transactivation potential, which corroborated that both proteins are functional HD-ZIP class I transcription factors.

### Two-rowed spikes have delayed lateral spikelet initiation and reduced growth

The size and fertility of lateral spikelets determine the row-type and intermedium-spike types in barley (Komatsuda et al., 2007; Ramsay et al., 2011; Youssef et al., 2017; Zwirek et al., 2019). To comprehend the differences of lateral and central spikelets in two-rowed barley, we tracked these spikelets from their early initiation until pollination in the two-rowed cv. Bowman. Barley spike development starts from the double ridge (DR) stage, in which spikelet ridges are subtended by leaf ridges (Fig. 2A). In the next stage, known as ‘triple mound’ (TM), the spikelet ridge differentiates into one central and two lateral spikelet meristems (CSM & LSM), in which the CSM develops as a bigger structure compared to the two LSMs (Fig. 2B). This marks the first difference between the central and lateral spikelets. Following the TM stage, the CSM continues to differentiate into various spikelet/floret organ primordia (glume, lemma, palea, stamen, pistil, and awn) (Fig. 2C-F). From the glume primordium stage, however, the LSM exhibits a delayed differentiation indicating the suppression of LSM (Fig. 2C). At the awn primordium stage (AP), the central spikelets have completed the differentiation of all spikelet/floret organs, while the laterals only achieved the differentiation of glume and lemma (Fig. 2F). We also compared the development of lateral spikelets between the two-rowed cv. Bowman and its near-isogenic six-rowed line BW-NIL(*vrs1*.*a*) (Druka et al., 2011). Close to the white anther stage (Kirby and Appleyard, 1984), the difference between the laterals of two- and six-rowed spikes became apparent (Fig. 3A-D). The six-rowed laterals possessed primordia for all spikelet/floret organs, whereas in two-rowed, laterals had retarded awn and pistil primordia (Fig. 3C & D). We also verified the divergence of lateral spikelet development in another pair of two- (cv. Bonus) and six-rowed (*hex-v*.*3, vrs1* deletion mutant) barleys (Supplementary Fig. 5). To fathom the sterility of lateral spikelets, we compared the histology of pistil and anther growth in Bowman and its *vrs1*.*a* mutant [BW-NIL(*vrs1*.a)] from Waddington stage 4.5 (W4.5, awn primordium stage) to W10.0 (pollination) (Supplementary Fig. 6&7). The delayed differentiation of lateral spikelets observed during the spikelet initiation stages (TM to AP) continued in the growth stages of the reproductive organs (Fig. 3E-L). Anthers of two-rowed lateral spikelets showed an impeded differentiation compared to the anthers of other spikelets (Supplementary Fig. 6, A3-J3). However, the central spikelet anthers of two- (Supplementary Fig. 6, A1-J1) and six-rowed (Supplementary Fig. 6, A2-J2) exhibited an advanced progression rate across the stages. Notably, the six-rowed lateral anther (Supplementary Fig. 6, A4-J4) followed a differentiation rate between the two- and six-rowed centrals as well as the two-rowed laterals, indicating that there are additional suppressors of lateral spikelet development besides HvHOX1. Moreover, anthers of the two-rowed lateral spikelets stopped differentiation at W7.5 (Supplementary Fig. 6, E3), followed by tissue disintegration in the subsequent stages (Supplementary Fig. 6, E3 to J3). In contrast, all other anthers continued their growth towards pollination (Supplementary Fig. 6). A similar delay of differentiation and disintegration of tissues was also observed in the pistil of two-rowed laterals at W7.5 (Supplementary Fig. 7, C5). Concertedly, these results substantiate that two-rowed spikes have delayed lateral spikelet initiation and suppressed growth compared to their central and all the spikelets of six-rowed spikes. Eventually, the reproductive organs of the lateral spikelets in two-rowed cv. Bowman abort during the later growth phase.

**Figure 2:**
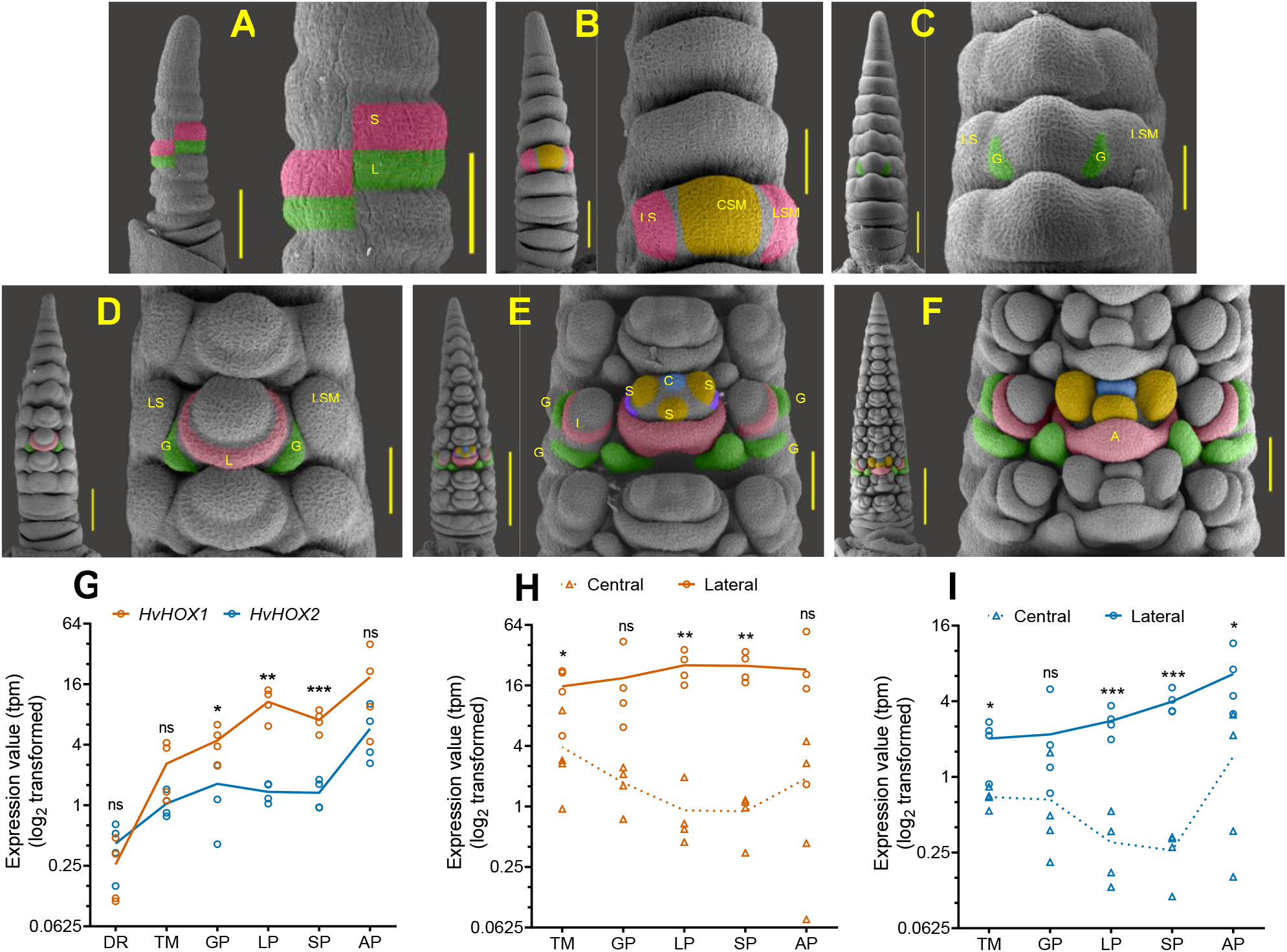
Two-rowed spikes have delayed lateral spikelet initiation and reduced growth. Early spike developmental stages of a two-rowed cultivar Bowman are shown from A to F. Double ridge (DR) is shown in A, in which the spikelet ridge (SR) is subtended by a leaf ridge (LR). The SR differentiates into one central and two lateral spikelet meristems (CSM & LSM) at triple mound stage (TM), which is displayed in B. Panel C discloses the appearance of two glume primordia (GP) from the CSM, while the two LSMs do not show any sign of differentiation. Subsequently, the CSM further differentiates and forms a lemma primordium (LP), which is shown in D. Two GP and a sign of LP initiation from the LSM can be seen in the panel E; the CSM initiated three stamen primordia along with a sign of carpel primordia development at this stage. At awn primordium stage (AP), F, the CSM completed the formation of all floral organ primordia (including the carpel), and the AP initiates from the medial point of the LP. However, the laterals are found only with two GP and a LP. Panels G, H, and I depicts the expression pattern of *HvHOX1* and *HvHOX2* genes in the whole spikes of DR to AP stages, *HvHOX1* and *HvHOX2* in the central and lateral spikelets of TM to AP stages, respectively. *HvHOX1* expresses higher than *HvHOX2* in the whole spikes of GP, LP, and SP stages (G). Both the genes are expressed in the dissected central and lateral spikelets from TM to AP stages. Mean values of G-I are compared with the multiple Student’s *t-*test; *, **, ***, mean values are significantly different at 5, 1, and 0.1% probability levels; ns-not significantly different. Scale bar in panel A - whole spike 500 µm, magnified three nodes 100 µm; B & C-500 µm & 200 µm; D-500 µm & 100 µm; E & F-200 µm & 100 µm. W-Waddington scale.

**Figure 3:**
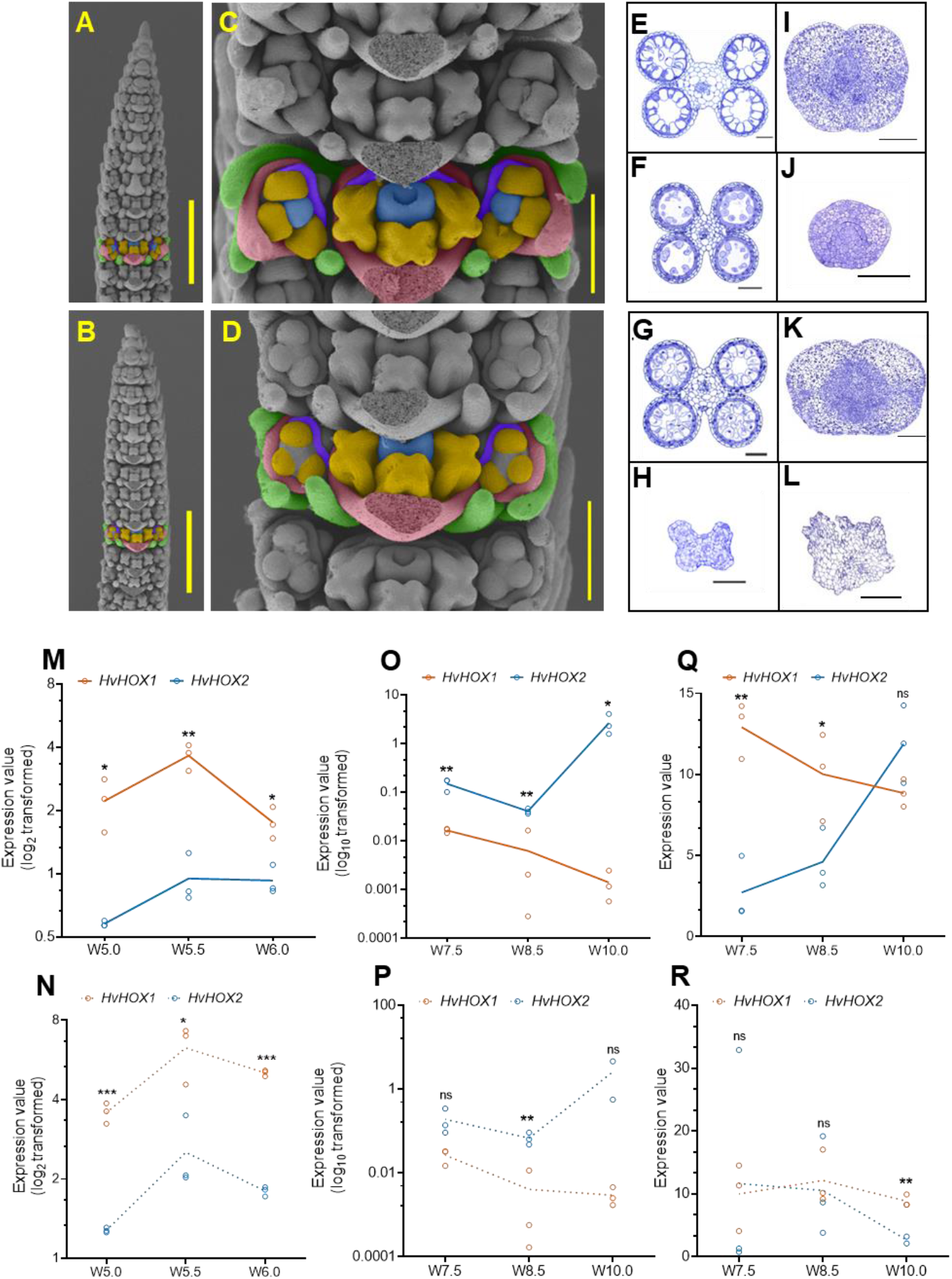
Two-rowed spikes have delayed and reduced lateral spikelet development compared to its central and six-rowed lateral spikelets. Images of panel A & C are the W5.5 stage inflorescence meristem of six-rowed mutant BW-NIL(*vrs1*.*a*), and B & D are from the two-rowed progenitor Bowman. Development of different organ primordia (AP, SP, CP, & GP) in central spikelets (yellow color) of two-rowed (D) and six-rowed (C) are visibly similar. Awn primordium (AP) and carpel primordium (CP) are formed only in lateral spikelets (blue color) of six-rowed (C) and not in two-rowed (D, marked with red arrow heads). Cross sections of anthers and carpels of BW-NIL(*vrs1*.*a*) and Bowman are shown in E-L. The W7.5 stage central spikelet anthers (E & G) and carpels (I & K) of BW-NIL(*vrs1*.*a*) and Bowman display normal development, while the lateral spikelet anther (H) and carpel (L) of Bowman show suppressed and aborted development. However, the lateral spikelet anther (F) and carpel (J) of BW-NIL(*vrs1*.*a*) seems developing normally but comparatively slower than its central spikelet organs. Expression of *HvHOX1* and *HvHOX2* genes in the whole spike (M & N), central spikelet (O & P), and lateral spikelet (Q & R) of Bowman and BW-NIL(*vrs1*.*a*) is shown, respectively. In whole spike of W5.0 to W6.0 stages, *HvHOX1* expressed greater than *HvHOX2* both in Bowman (M) and BW-NIL(*vrs1*.*a*) (N). Contrastingly, the *HvHOX2* showed stronger expression than *HvHOX1* in the Bowman central spikelets of W7.5 to W10.0 (O); however, in BW-NIL(*vrs1*.*a*), *HvHOX2*’s expression was higher only in W8.5. In the Bowman lateral spikelets of W7.5 to W10.0, *HvHOX1* and *HvHOX2* exhibited an anticyclic expression pattern, i.e., when *HvHOX1*’s expression dropped down from W7.5 to W10.0, *HvHOX2*’s expression started increasing. Mean values of M-R are compared with the multiple Student’s *t-*test; *, **, ***, mean values are significantly different at 5, 1, and 0.1% probability levels; ns-not significantly different. orange-stamen primordium; blue-carpel primordium; green: glume primordium; pink: lemma primordium; purple: palea primordium; W-Waddington scale. Scale bar, A&B, 800 µm; C&D, 200 µm; E-L is 100 µm.

### *HvHOX1* and *HvHOX2* have a contrasting dosage of expression during spikelet initiation and growth

We have taken the log_2_ transformed expression values of *HvHOX1* and *HvHOX2* from the Bowman RNA-seq spike atlas data (Thiel et al., 2021) and reanalyzed them to find their expression pattern across the spikelet initiation stages (Fig. 2G-I). In the tissue-unspecific (central and lateral combined or whole spike) transcript analysis, both genes showed a linear increase in expression along with the spikelet initiation stages (Fig. 2G). With the exception of the DR stage, *HvHOX1* generally displayed higher transcript levels than *HvHOX2* (TM to AP). This was particularly evident in glume primordium (GP), lemma primordium (LP), and stamen primordium (SP) stages (Fig. 2G). We then compared the tissue-specific expression patterns of these genes in central and lateral spikelets. Notably, both genes had higher levels of mRNA in the laterals than centrals at TM, LP, and stamen primordium (SP) stages (Fig. 2H & I). Then, we compared the expression level of these genes within the same tissues (central and lateral spikelets), in which *HvHOX1* showed significantly higher expression than *HvHOX2* in several stages (TM, LP, & SP) of lateral and at the SP stage of central spikelets (Supplementary fig. 8A & B). The high expression of *HvHOX1* in the laterals correlates with the delayed differentiation and suppression of the lateral spikelets (compared to the centrals) from the TM to AP stages in Bowman (Supplementary Fig. 8B). This reinforced the role of HvHOX1 as a negative regulator of lateral spikelet development in barley (Komatsuda et al., 2007; Sakuma et al., 2010; Sakuma et al., 2013). The presence of *HvHOX1* transcripts in central spikelets of two-rowed barleys, which are fertile and do not show any developmental disorder, poses a question that has yet to be solved (Komatsuda et al., 2007; Sakuma et al., 2010; Sakuma et al., 2013) (Fig. 2C-F, Supplementary Fig. 7&8).

Following the comparison on spikelet initiation stages, we explored expression levels of these genes also in the spikelet growth stages of Bowman and BW-NIL(*vrs1*.*a*) (non-functional *HvHOX1*) by doing a quantitative real-time (qRT) PCR with tissue-unspecific (W5.0, W5.5, & W6.0) and tissue-specific (W7.5, W8.5, W10.0) samples (Fig. 3M-R). Also, in these later stages of development, *HvHOX1* exhibited significantly higher expression than *HvHOX2* in the whole spike at W5.0, W5.5, and W6.0, both in Bowman and BW-NIL(*vrs1*.*a*) (Fig. 3M & N). Intriguingly, *HvHOX1* displayed a reduced expression trend both in the central and lateral spikelets of Bowman from W7.5 to W10.0 (Fig. 3O & Q). Contrastingly, *HvHOX2* had an increasing trend of expression in these two tissues of Bowman (Fig. 3O & Q). More importantly, *HvHOX2* showed greater expression than *HvHOX1* in the centrals (Fig. 3O), whereas in the laterals, *HvHOX1* had a superior level of expression in the first two stages (W7.5 and W8.5), followed by the increase of *HvHOX2* at W10.0 (Fig. 3Q). Crucially, the transcript levels of *HvHOX1* were gradually decreased from W7.5, while *HvHOX2* levels increased. Similar to the Bowman centrals, *HvHOX2* showed a higher trend of expression in BW-NIL(*vrs1*.*a*) central spikelets. However, the expression patterns of these two genes were different in BW-NIL(*vrs1*.*a*) lateral spikelets compared to Bowman (Fig. 3P & R). The antagonistic expression patterns of *HvHOX1* and *HvHOX2*, i.e., when *HvHOX2* expression goes up, *HvHOX1* expression turns down, suggests that these two genes might act anti-cyclic during the later growth stages. Based on this observation and the higher expression of *HvHOX2* (W7.5 to W10.0) in the Bowman central spikelets (Fig. 3O) that show no developmental and growth aberration, we hypothesized that overexpression of *HvHOX2* might promote spikelet development by acting as a positive regulator of spikelet development.

### Promoters of *HvHOX1* and *HvHOX2* share similar spatiotemporal expression patterns during spike growth stages

The expression studies of *HvHOX1* and *HvHOX2* (Fig. 2G-I & 3M-R) exemplified that these genes have similar temporal expression during the spikelet initiation and growth stages though at different amplitudes. Additionally, their central- and lateral-specific transcript levels indicated that they might also share spatial boundaries across the initiation and growth stages. To verify their spatial co-localization and similar temporal expression patterns, promoters (*HvHOX2*-1929 bp; *HvHOX1*-991 bp) of these genes were fused with a synthetic *GFP* (*GFP*) coding sequence and transformed into the two-rowed cv. Golden Promise. Five and eight independent transgenic events showed GFP accumulation in the T_0_ generation for *HvHOX1*, and *HvHOX2* GFP constructs, respectively. Three independent events from both the constructs were selected, and their GFP accumulation was confirmed until T_2_ generation. As expected, we found that promoter activity of these genes in identical tissues like the base of the central spikelet’s carpel (Fig. 4A & D), the tapetal layer of the central spikelet’s anther (Fig. 4B E), and rudimentary lateral anthers (Fig. 4C & F) at W8.5 stage. Collectively, the tissue-specific expression analysis and the promoter activity in the transgenic plants suggested that *HvHOX1* and *HvHOX2* might have similar spatiotemporal expression patterns during spikelet growth stages.

**Figure 4.**
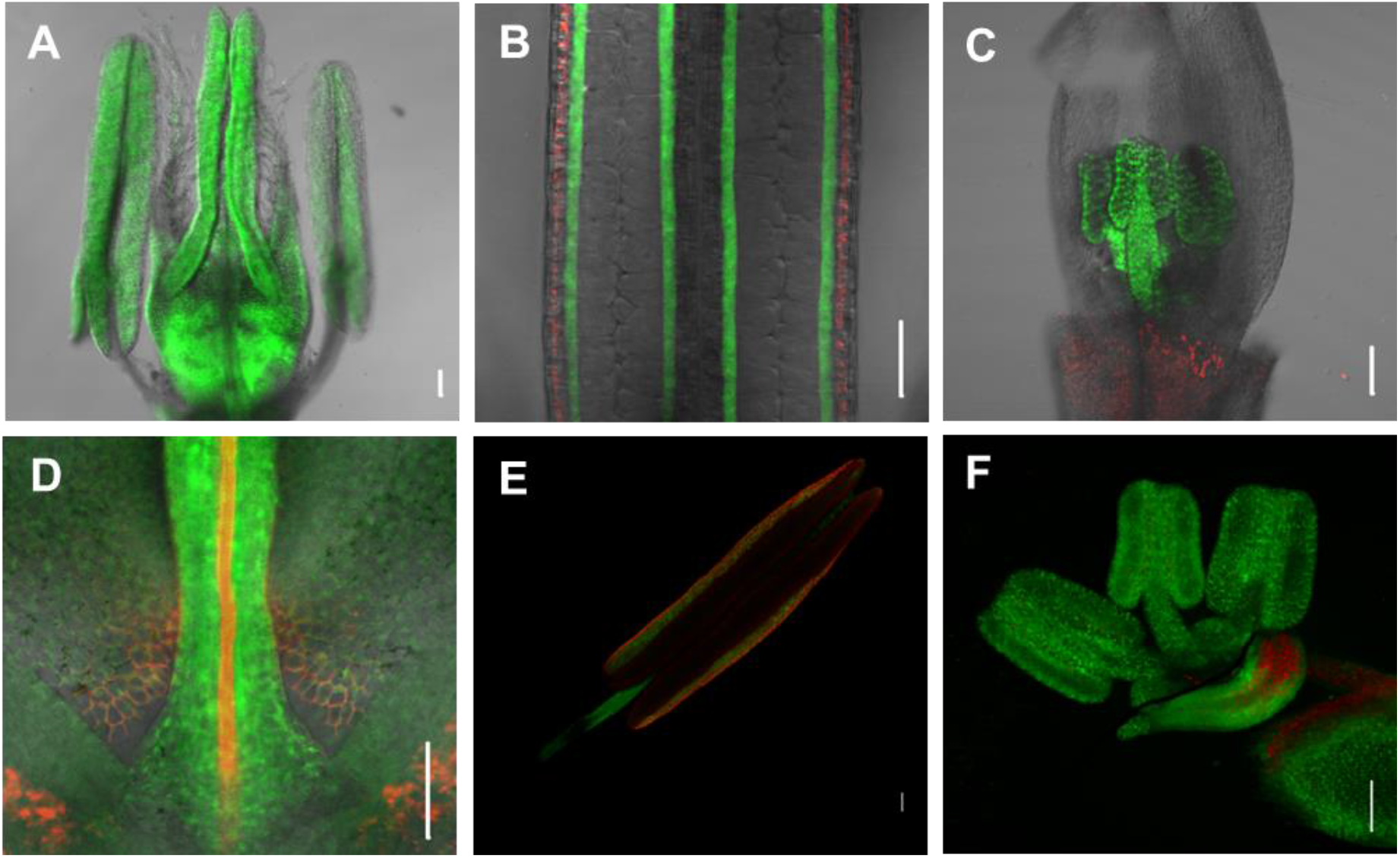
*HvHOX1* and *HvHOX2* have similar spatiotemporal expression pattern during spike growth and development. *HvHOX2* promoter activity (GFP expression) in central spikelet’s stamen and carpel (A), tapetum of central spikelet’s stamen (B) and lateral spikelets’ stamen (C) at W8.5 is shown. Similarly, *HvHOX1* promoter activity in central spikelets’ carpel (D), tapetum of central spikelet’s stamen (E) and lateral spikelet’s stamen (F) is shown. Green color - GFP fluorescence; red color-chlorophyll autofluorescence. Scale bar 100 µm. W-Waddington scale.

### *HvHOX1* has a unique co-expression module apart from a shared module with *HvHOX2* during spike development

In an effort to predict the role of *HvHOX1* and *HvHOX2* genes, we constructed their co-expression signatures from the transcript profiles across six spikelet initiation and growth stages (W2.5, W3.0, W4.5, W6.5, W7.5, and W8.5) in Bowman. We found twenty co-expression modules from a set of 7,520 genes that showed a dynamic expression profile (Fig. 5A & B). *HvHOX1* and *HvHOX2* genes clustered together in one module (Figure 5A; red) along with 4,213 genes. A weighted gene co-expression network analysis (WGCNA) revealed that *HvHOX1* shares one part of its co-expression module (Fig. 5C, shown in blue, 16 genes) with *HvHOX2*, while *HvHOX1* has unique co-expressed signatures (Fig. 5C, shown in orange, 39 genes). Most importantly, *HvHOX2* is one of the co-expressed genes within the *HvHOX1* module (Fig. 5C). In other words, both genes share a similar expression signature across spike development. This supports our previous transcript and GFP analyses and suggests that these genes have similar spatiotemporal expression patterns. Furthermore, hierarchical clustering (HCL), divided the genes in the shared module (Supplementary fig. 9A; blue) into two sub-clusters based on their expression in central and lateral spikelets, but not the unique *HvHOX1* co-expressed module (Supplementary Fig. 10B). This indicates that HvHOX1 may play a specific role in the lateral spikelets, while HvHOX2 probably has a different function from HvHOX1. Interestingly, the shared module was enriched with genes [e.g. *AGAMOUS* (*AG*), *SUPPRESSOR OF OVEREXPRESSION OF CONSTANS 1* (*SOC1*), *ENOLASE 1* (*ENO1*), and *AUXIN F-BOX PROTEIN 5* (*AFB5*)] associated with flower development, promotion of flowering, carpel and stamen identity, auxin signaling, transcription and nitrate assimilation (Covington and Harmer, 2007; Dreni and Kater, 2014; Hyun et al., 2016; Gaufichon et al., 2017). The *HvHOX1* unique co-expressed module, on the other hand, was enriched in genes [such as *BREVIPEDICELLUS 1* (*BP1*), *WRKY 12, NOVEL PLANT SNARE 11* (*NPSN11*), *FORMIN HOMOLOGY 14* (*AFH14*), *LONELY GUY 3* (*LOG3*), and *G PROTEIN ALPHA SUBUNIT 1* (*GPA1*)] that are predicted to be involved in inflorescence architecture, flower development, ABA response, cell division, cell communication, senescence, and cell death (Li et al., 2010; Tokunaga et al., 2012; Zhao et al., 2015; Li et al., 2016; Chakraborty et al., 2019; Wu et al., 2020) (Supplementary Table 4).

**Figure 5.**
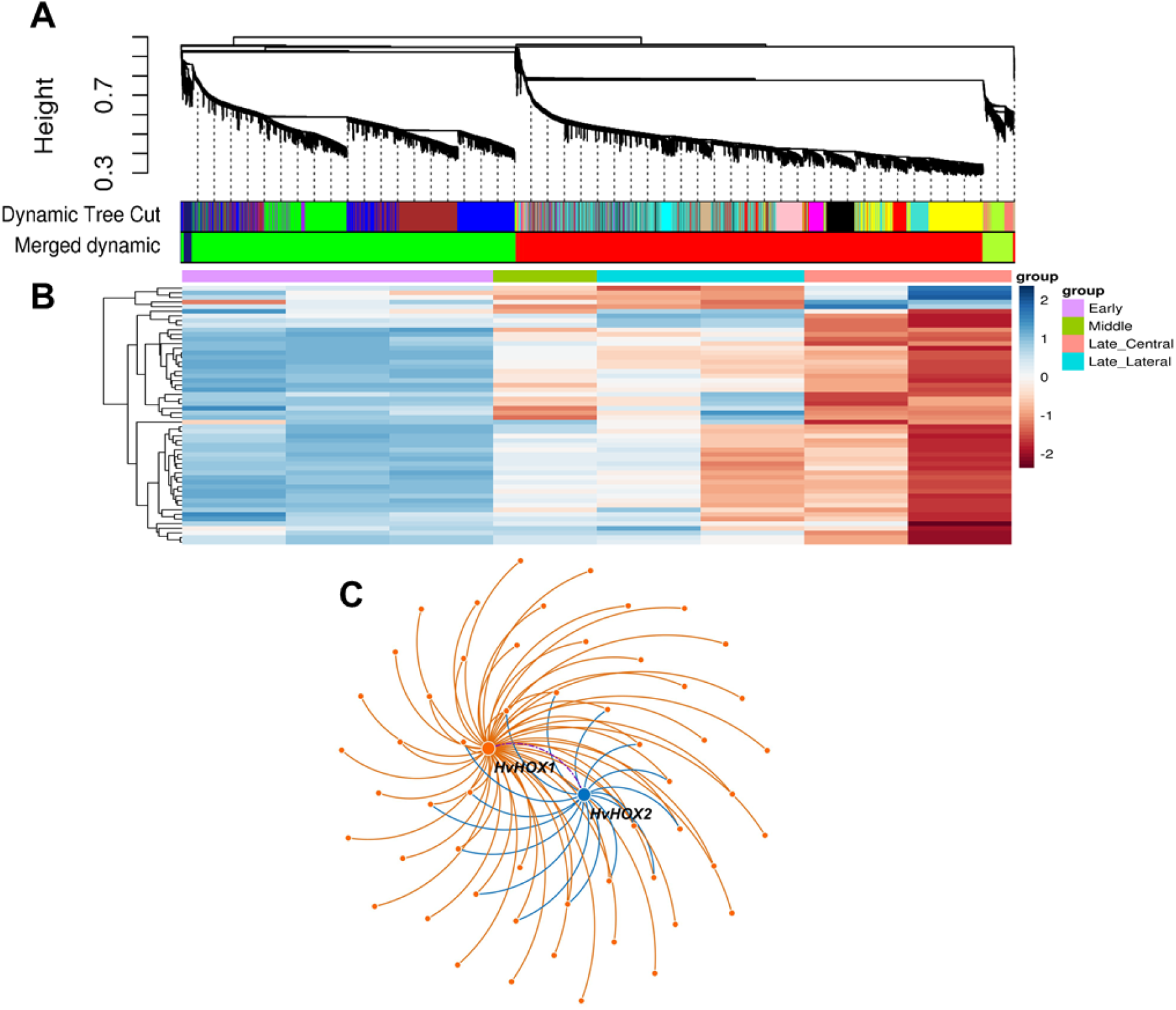
Dendrogram from gene co-expression network analysis of two-rowed cv. Bowman spike tissues. Modules of the co-expressed genes were assigned colors, shown by the horizontal bars below the dendrogram. Merged modules are shown under the dynamic module profile (A). Expression heat map of the red module is shown in (B) and the coexpressed gene clusters of *HvHOX1* and *HvHOX2* are shown in (C).

### *HvHOX2* might be a dispensable gene during barley spikelet development

To understand the function of *HvHOX2*, we developed *Hvhox2* mutants by using RNA-guided Cas9 endonucleases (RGEN). A guide RNA was designed for the conserved homeodomain region shared by *HvHOX1* and *HvHOX2* for site-directed mutagenesis of both genes (Fig. 6A). We created the mutants in the two-rowed cv. Golden Promise, via stable transformation, using respective constructs and identified the independent events BG724_E02 and BG724_E07 bearing different insertions and/or deletions by sequencing their target regions. Among the progenies of these primary (heterozygous/chimeric) mutants, wild-type (T-DNA-free, non-mutant) plants, as well as single and double mutants for both genes, were selected (Fig. 6A). For *HvHox1*, the two mutants, BG724_E02 and BG724_E07, had one and eight nucleotides deletions, respectively, in the target regions (Fig. 6A), which created two frame shifted mutant HvHOX1 proteins (Supplementary Fig. 10A&B). With regards to *Hvhox2*, the BG724_E02 event had seven nucleotides addition and four nucleotides deletion (Fig. 6A), which resulted in a mutant HvHOX2 protein that had one amino acid addition and one amino acid exchange in the first HD (Supplementary Fig. 10C). Similar to the *Hvhox1* BG724_E02 mutant, *Hvhox2* BG724_E07 mutant possessed one nucleotide deletion (Fig. 6A) and formed a frame shifted protein (Supplementary Fig. 10D). The spikelet development of these plants was compared at W4.5 and after spike maturity. It was found that the central and lateral spikelets of the *Hvhox2* mutants (Fig. 6C, Supplementary Fig. 11B) displayed a similar stage of differentiation at W4.5 as in the spikes of wild-type plants (Fig. 6B, Supplementary Fig. 11A). Analogous to the pattern of spikelet differentiation, the matured spikes of *Hvhox2* mutants (Fig. 6G, Supplementary Fig. 11F) possessed smaller (compared to the centrals) and sterile lateral spikelets like in spikes of wild-type plants (Fig. 6F, Supplementary Fig. 11E), implying that *HvHOX2* might neither promote nor suppress spikelet primordia differentiation and growth. However, *Hvhox1* single (Fig. 6D, Supplementary Fig. 11C) and double mutants (*Hvhox1/Hvhox2*) (Fig. 6E, Supplementary Fig. 11D) exhibited advanced lateral spikelet differentiation compared to wild-type plants (Fig. 6B, Supplementary Fig. 11A) and *Hvhox2* mutants (Fig. 6C, Supplementary Fig. 11B). Interestingly, the lateral spikelet differentiation of *Hvhox1* (Fig. 6D, Supplementary Fig. 11C) and double mutants (*Hvhox1/Hvhox2*) (Fig. 6E, Supplementary Fig. 11D) were at a similar stage at W4.5, which reiterated the fact that *HvHOX1* is suppressing lateral spikelet development in two-rowed spikes, irrespective of the *HvHOX2* function. As expected, spikes of *Hvhox1* single (Fig. 6H, Supplementary Fig. 11G) and double mutant (*Hvhox1/Hvhox2*) (Fig. 6I, Supplementary Fig. 11H) had bigger and fertile spikelets (grains) like six-rowed barley. We explored *Hvhox2* mutants by screening its coding sequence in 5500 second-generation (M_2_) TILLING (Targeting Induced Local Lesions in Genomes) mutant lines of cv. Barke (Gottwald et al., 2009), and found only four mutations. Among these, three were synonymous, and one was non-synonymous (P197S, line 11869) nucleotide substitutions (Supplementary Fig. 12). Interestingly, the mutant line 11869 did not show aberrations during spike development and growth in the M_3_ generation, which supported our RGEN *Hvhox2* mutants. Taken together, our RGEN mutant analyses suggest that *HvHOX2*, at its native expression level, appears dispensable for barley spikelet development.

**Figure 6:**
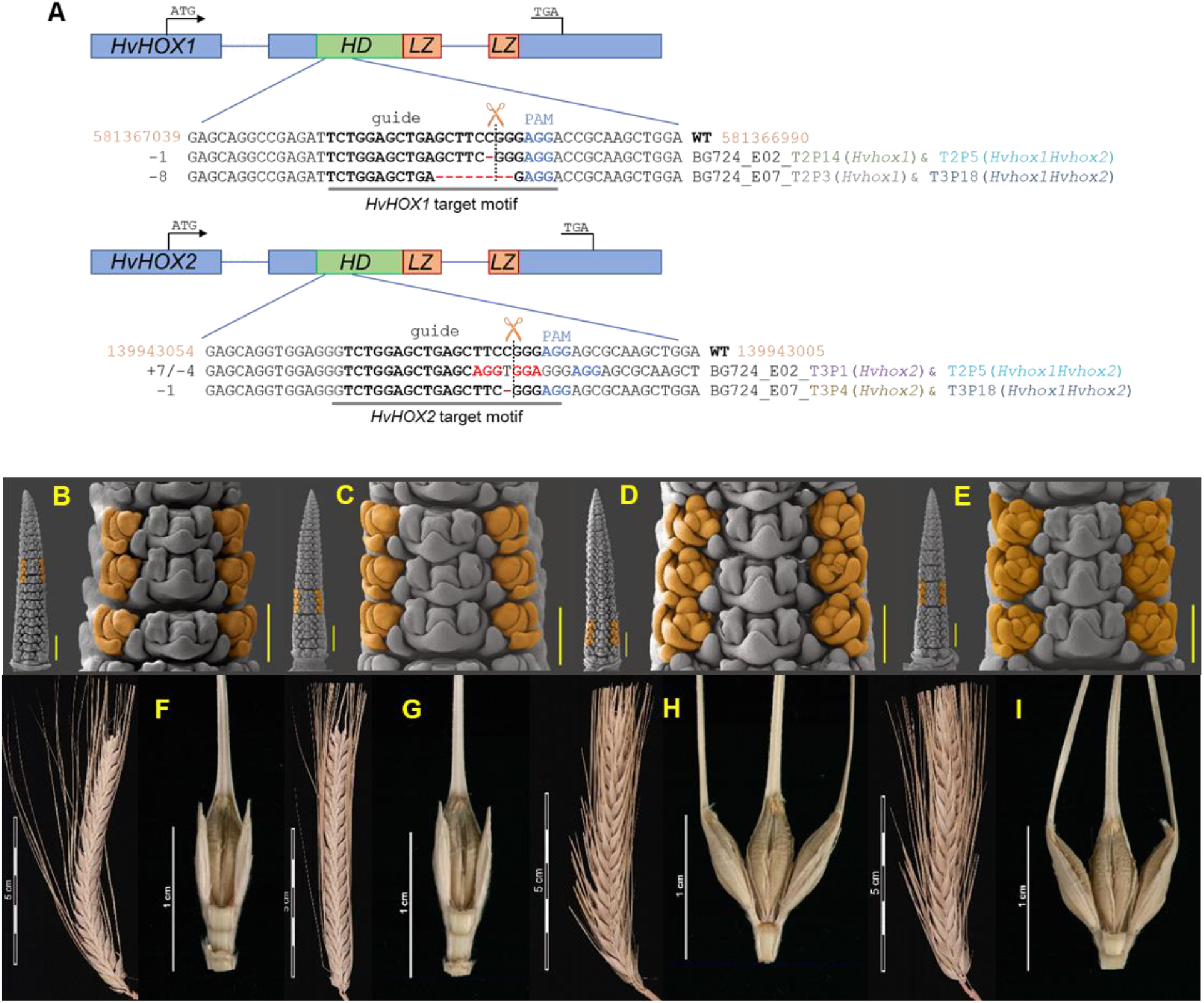
*HvHOX2* gene might be a team player in barley spikelet development. Figure A graphically shows the guide sequence with the Protospacer Adjacent Motif (PAM) and a putative cutting site, used to generate the single and double mutants of *HvHOX1* and *HvHOX2* genes, by using Cas9 endonuclease. Figures B & F are from an azygous plant and C-D & G-I are the representative images of the BG724-E07 mutants. Figures B-E compare the lateral spikelet development of wild-type, single and double mutants of *HvHOX1* and *HvHOX2* genes. At W4.5, the lateral spikelet primordia of *HvHOX2* mutant (C) is at similar developmental stage with the wild-type (B) by having differentiated primordia for only glume and lemma. However, the mutant of *HvHOX1* (D) and double mutant of *HvHOX1* and *HvHOX2* (E) displayed an advanced lateral spikelet development with the well differentiated primordia for glume, lemma, stamen, and carpel. The matured lateral spikelets of *HvHOX2* mutant (G) and wild-type (F) are sterile and smaller compared to the fertile lateral spikelets that form grains of *HvHOX1* mutant (H) and *HvHOX1* and *HvHOX2* double mutant (I). Scale bar-whole spikes in B, C, D, & E, 500 µm; magnified three nodes, 200 µm; HD-Homeodomain; LZ-Leucine Zipper.

### Overexpression of *HvHOX2* can promote lateral spikelet development

Our qRT expression study conducted during spike growth stages revealed that higher transcript levels of *HvHOX2* than *HvHOX1* in central spikelets might be associated with the proper development of those spikelets in two-rowed barley (Fig. 3D, G, K & O). To validate this ‘*HvHOX2*-dosage’-hypothesis, we tagged the *HvHOX1* promoter (991 bp – also used for assessing the spatiotemporal activity of *HvHOX1* promoter) with the coding sequence of *HvHOX2* and used these constructs to create transgenic plants of cv. Golden Promise. We used the *HvHOX1* promoter because *HvHOX1* expresses higher in the lateral spikelets (Supplementary Fig. 8B, Fig. 3Q), so this promoter might increase the transcript levels of *HvHOX2* in the lateral spikelets of transgenic plants. As a result, the smaller and sterile lateral spikelets might be restored to fertile and bigger spikelets. Eight independent transgene-positive events were selected and screened for the restored lateral spikelets. Two events, E189 (at T_2_) and E541 (at T_1_), showed partial promotion of lateral spikelets compared to a wild-type control plant E511 (Fig. 7). The spikes of the two events displayed an advanced lateral spikelet differentiation at W4.5 compared to the spike of wild-type (E511) plants (Fig. 7A-C). Interestingly, the lateral spikelets of both the events had a quantitative difference in development, in which E189 showed a mild promotion, while E541 possessed a bit stronger improvement compared to the spikes of control plants (Fig. 7B & C). The matured spikes of E189 and E541 had partially restored lateral spikelets that are bigger and occasionally developed small awns in contrast to the spikes of control plants (Fig. 7D-F). The matured lateral spikelets of E189 were smaller than E541, which followed the similar pattern of developmental differentiation observed during spikelet differentiation (Fig. 7B&C and E&F). Then, we quantified the transcripts of *HvHOX1, HvHOX2*, and *HvHOX2-T* (*HvHOX1pro::HvHOX2*) in W6.5 (tissue-unspecific) (Fig. 7G) and W8.0 (tissue-specific) (Fig. 7H&I). It revealed that both the events (E189 & E541) had *HvHOX2-T* transcripts in the two stages and tissues analyzed (Fig. 7G-I). Most importantly, there was no difference in the expression levels of *HvHOX1* and *HvHOX2* genes in the lateral spikelets of transgenic events compared to the azygous plant (Fig. 7I). However, we found a significant reduction of *HvHOX1* transcripts in the central spikelets (Fig. 7H). We also observed that event E189 had a four-fold higher expression of *HvHOX2-T* than E541 at W6.5 (Fig. 7G), which was similarly seen in the lateral spikelets at W8.0, where E189 had 1.6-fold higher expression than E541 (Fig. 7I). We hypothesize that this difference in expression is mainly due to the developmental disparity between E189 and E541 (Fig. 7B & C). Thus, our overexpression study supports the idea that increasing the dosage of *HvHOX2* transcripts promotes lateral spikelet development in two-rowed barley.

**Figure 7:**
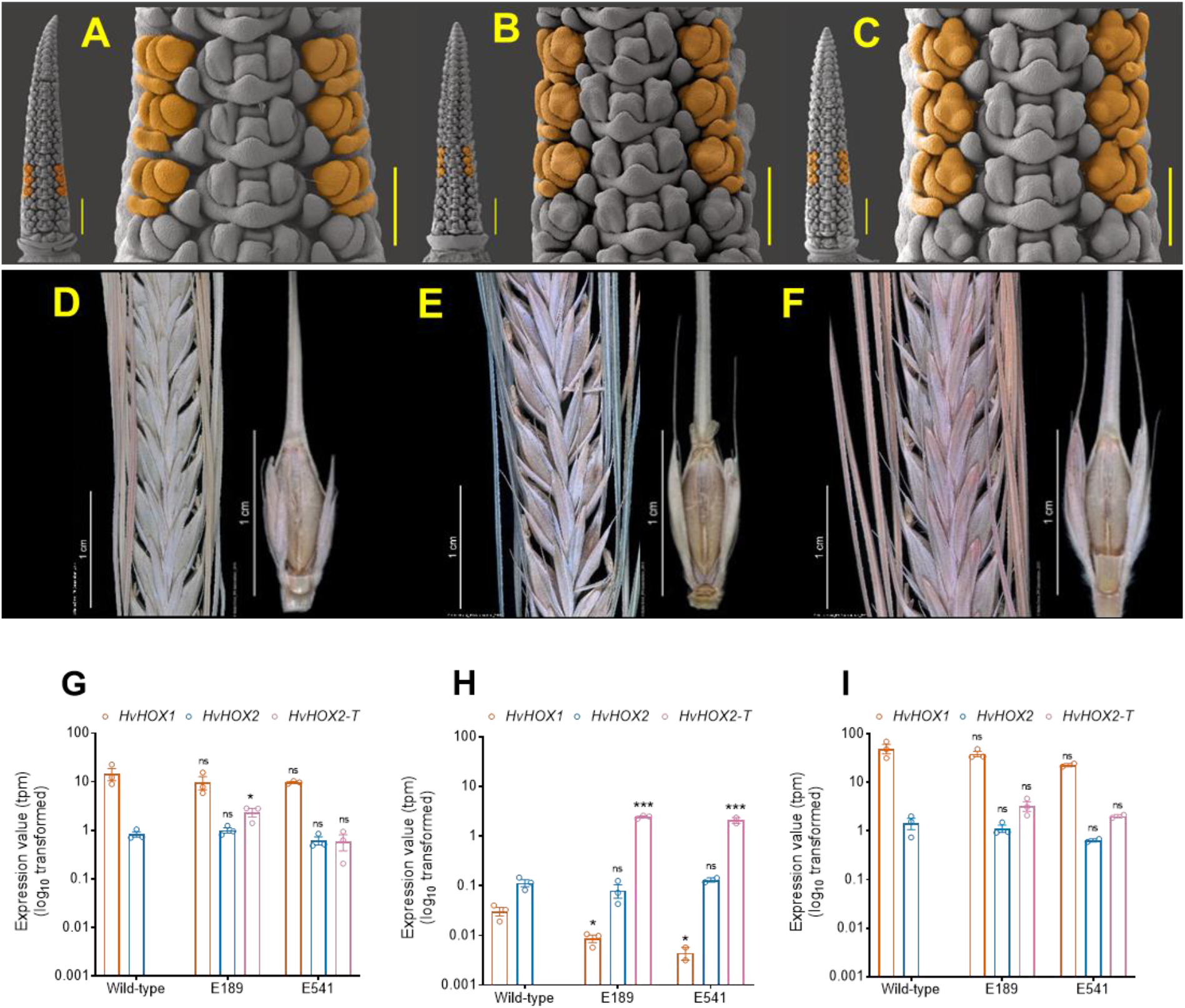
Overexpression of *HvHOX2* partially restored the lateral spikelet development in two-rowed barley. A comparison of wild-type (A) and two transgenic plants E189 (B) E541 (C) lateral spikelet primordia differentiation is displayed. The lateral spikelet primordia of transgenic plants E189 and E541 exhibited an advanced development compared to the wild-type at W4.5. At this stage, the wild-type laterals are found only with the differentiated glume and lemma primordia, while both the transgenic plants already initiated three stamen primordia along with the glume and lemma primordia. Matured spikes and triple spikelets of wild-type (D), E189 (E), and E541 (F) are shown. The lateral spikelets of both the transgenic plants are bigger compared to the wild-type and occasionally found with short awns. Quantification of endogenous *HvHOX1* and *HvHOX2* and transgenic *HvHOX2-T* expression performed in a wild-type and two independent transgenic plants’ (E189 & E541) whole spike at W6.5 (G), central spikelet (H) and lateral spikelets (I) at W8.0 is shown. The overexpression of transgenic *HvHOX2-T* did not greatly change the endogenous *HvHOX1* and *HvHOX2* expression in the whole spike at W6.5 (G) and lateral spikelets of W8.0 (I). However, in the central spikelets of W8.0, the transgenic *HvHOX2-T* expression drastically lowered the *HvHOX1* expression (H). In G, H & I, the mean values of *HvHOX1* from the transgenic plants were compared to the wild-type. Similarly, both the endogenous and transgenic *HvHOX2* of transgenic plants were compared with the wild-type *HvHOX2* expression. Mean values of G-I are compared with the multiple Student’s *t-*test; *, ***, mean values are significantly different at 5 and 0.1% probability levels. In A, B, & C, the scale bars of whole spike images represent 500 µm, and in magnified three node images they represent 200 µm. W-Waddington scale.

## Discussion

### HvHOX1 and HvHOX2, two functional HD-ZIP class I transcription factors, may act antagonistically to each other

Based on the sequence similarity of *HvHOX2* to its orthologs in grass species, it was proposed that HvHOX2 might have a similar molecular role in the Poaceae (Sakuma et al., 2010). However, *HvHOX1*, specific to the Triticeae tribe, showed a very high sequence variation, at least in barley (Komatsuda et al., 2007; Saisho et al., 2009; Casas et al., 2018). It was hypothesized that *HvHOX1* and *HvHOX2* are duplicated genes, in which *HvHOX2* might be retaining the ancestral sequence and promotion of development, while *HvHOX1* became neofunctionalized as a suppressor of lateral spikelets (Sakuma et al., 2010; Sakuma et al., 2013). Our nucleotide diversity study also supports this postulation, as we found a higher nucleotide diversity for *HvHOX1* than *HvHOX2* (Supplementary Fig. 1, Supplementary Table 3). Despite a few amino acid changes between HvHOX1 and HvHOX2 proteins (Supplementary Fig. 3), both of them can bind to their HD-ZIP class I-specific cis-element, make dimers, and transactivate their downstream genes (Fig. 1, Supplementary Fig. 3&4), thus confirming that both are functional HD-ZIP class I TFs. Also, our expression studies suggested that both the genes have similar spatiotemporal expression patterns during spikelet initiation (Fig. 2) and growth stages (Fig. 3 & 4) that could facilitate the interaction between them. Similarly, our gene co-expression network (GCN) analysis revealed that most likely, these genes are sharing similar gene networks, as they fall into the same cluster of co-expressed genes and share a common network of genes (Fig. 5). This finding reaffirms the hypothesis that both genes might have originated from a common ancestral gene (Sakuma et al., 2010; Sakuma et al., 2013). Crucially, *HvHOX1* has a unique network of genes (Fig. 5C) that are highly expressed in lateral spikelets (Supplementary Fig. 9B) and are enriched with genes that are involved in the suppression of development and exert cell death (Supplementary Table 4) (Thirulogachandar et al., 2017). It also corroborates that *HvHOX1* might have acquired a new role as a suppressor of lateral spikelets in barley. Contrastingly, the genes in the shared network of *HvHOX1* and *HvHOX2* are expressed both in the central and lateral spikelets, and they are predicted to function towards the promotion of development and flowering. It also suggests that along with the suppressors, there might also be some promoters that are highly expressed in the lateral spikelets. Presumably, this is the first insight into the antagonistic behavior of these two genes during barley spikelet development.

Additionally, our analyses of differentially expressed genes between Bowman and BW-NIL(*vrs1*.*a*) (Supplementary Fig. 13&14) and wild-type and *HvHOX2* overexpressing transgenic plants (Supplementary Fig. 15) pointed out that *HvHOX1* and *HvHOX2* might work antagonistically to each other during spikelet development. There are many examples in plants in which homologous/paralogous genes are antagonists. In Arabidopsis, WRKY12 and WRKY13 oppositely modulate flowering time under SD conditions; WRKY12 promotes flowering, whereas WRKY13 delays this process (Li et al., 2016). Likewise, TERMINAL FLOWER 1 (TFL1) and FLOWERING LOCUS T (FT) are homologous PEBP class proteins, which are antagonistic to each other; TFL1 being a repressor and FT an activator of flowering (Hanzawa et al., 2005). Another example is the closely related MADS-box proteins SHORT VEGETATIVE PHASE (SVP) and AGAMOUS-LIKE 24 (AGL24), which perform opposite roles during the floral transition, acting as repressor and promotor of flowering, respectively (Hartmann et al., 2000; Yu et al., 2002; Michaels et al., 2003; Lee et al., 2007). Recently, in rice, it was found that *Teosinte branched 2* (*Tb2*) counteracts with its paralog *Tb1* to influence tiller number(Lyu et al., 2020). All these instances corroborate that gene duplication events followed by neofunctionalization might generate homologous/paralogous genes that can act antagonistically to each other and modulate specific developmental pathways. To understand the evolutionary importance of these genes, a new sub-category under neofunctionalization might be necessary for which we propose to group them as ‘antifunctionalized’ homologs. Thus, our studies suggest that the paralogous HD-ZIP class I transcription factors, HvHOX1, and HvHOX2 are antifunctionalized and may act against each other during barley spikelet development.

### Dosage of *HvHOX1* and *HvHOX2* transcripts influence spikelet development

HvHOX1 was previously proposed as a negative regulator of lateral spikelet, specifically pistil/carpel development, in barley (Komatsuda et al., 2007; Sakuma et al., 2010; Sakuma et al., 2013). We found evidence supporting the claim that from the initiation of TM (Fig. 2G & H) to W8.5 (Fig. 3M & Q), *HvHOX1* transcripts are enriched in the lateral spikelets of two-rowed barley. This correlated well with the delayed meristem differentiation (Fig. 2B-F) and anther and carpel development within the lateral spikelets (Fig. 3E-L, Supplementary Fig. 6&7). More importantly, the abortion of lateral spikelets’ anther and pistil/carpel at W7.5 (Fig. 3E-L, Supplementary Fig. 6&7), and the gradual reduction of *HvHOX1* expression in lateral spikelets from W7.5 (Fig. 3Q), reaffirm that *HvHOX1* is highly expressed in the reproductive organs of lateral spikelets. We also identified *HvHOX1* transcripts in central spikelets during early and late spikelet development (Fig. 2H; Fig. 3O & Q). However, we observed no disorder during spikelet differentiation (Fig. 2B-F, 3C & D) or growth of reproductive organs (Fig. 3E-L, Supplementary Fig. 6&7) in two-rowed barley. Also, previous studies did not report any developmental irregularities in central spikelets of two-rowed barley (Komatsuda et al., 2007; Sakuma et al., 2010; Sakuma et al., 2013; Zwirek et al., 2019). We, therefore, hypothesized that this could be due to a lower dosage of *HvHOX1* transcripts (compared to the laterals) (Fig. 2H & 3O) and some more positive regulators, which act antagonistically to *HvHOX1* in central spikelets. It led us to examine the expression of *HvHOX2* - a paralog of *HvHOX1*, which was proposed to be promoting the development in barley (Sakuma et al., 2010; Sakuma et al., 2013). A similar (i.e., non-significant) level of *HvHOX2* transcripts as *HvHOX1* during the early spikelet differentiation stages (Supplementary Fig. 8A) (except at SP stage), and a higher dosage of *HvHOX2* transcripts in the central spikelets across the growth of reproductive organs (Fig. 3O) supports the claim of a promoting HvHOX2 function. Furthermore, we recognized an anti-cyclic expression pattern between these two genes during the growth stages (Fig. 3Q) and binding of HvHOX1 protein on *HvHOX2* promoter and vice versa (Supplementary Fig. 4) indicating that these genes influence the expression pattern of each other. A similar expression pattern of these two genes had already been reported in other two-rowed barleys(Sakuma et al., 2013).

Our RGEN mutant study suggests that HvHOX2 is rather dispensable during barley spikelet development because the two *Hvhox2* mutants retained a canonical spikelet development in laterals and centrals of wild-type plants (Fig. 6, Supplementary Fig. 10). Interestingly, ubiquitous overexpression of orthologous *HOX2* genes in wheat(Wang et al., 2017) and rice(Shao et al., 2018) reduced the inflorescence length and complexity. However, when *HvHOX2* transcripts were increased transgenically, HvHOX2 can restore and promote barley lateral spikelet development in a dosage-dependent manner (Fig. 7). A significant reduction of *HvHOX1* transcripts in the central spikelets of *HvHOX2* overexpression mutants (Fig. 7H) reinstated that these two genes can influence each other’s expression level. We also observed a reduction of *HvHOX1* transcripts in the lateral spikelets of *HvHOX2* overexpression plants. Specifically, 3.5 and 6.9 times (mean transcript values) of reduction in transcripts were identified in the *HvHOX2* overexpressing plants E189 and E541, respectively; however, the declines were not statistically significant (Fig. 7I). We hypothesize that this could be due to the solid lateral-specific expression of *HvHOX1*, which is under the control of VRS4 (HvRA2) and VRS3 – two upstream regulators of *HvHOX1* (Koppolu et al., 2013; Sakuma et al., 2013; Bull et al., 2017; van Esse et al., 2017). The reduction level of *HvHOX1* transcripts and the degree of lateral spikelet promotion in the two *HvHOX2* overexpression events indicated that HvHOX1 regulates lateral spikelet development based on the dosage of its expression, which was also shown previously (Sakuma et al., 2013). Taken together, our expression and transgenic studies suggest that the transcript levels of *HvHOX1* and *HvHOX2* influence lateral spikelet development in two-rowed barley in a dosage-dependent fashion.

## Methods

### Plant materials and their growth conditions

Barley cultivars, Bonus, Bowman, and Golden Promise, were used in this study as two-rowed representatives and induced mutant *hex-v*.*3* (progenitor cv. Bonus), cultivar Morex and Bowman backcross-derived line BW-NIL(*vrs1*.*a*) / BW 898 (Druka et al., 2011) were used as six-rowed representatives. Wild species of *Hordeum* were obtained from Dr. Roland von Bothmer, Swedish University of Agricultural Sciences, Alnarp, Sweden (Supplementary table 2). *Arabidopsis thaliana* Col-0 plants were used for protoplast isolations and grown on a 1:3 vermiculite: soil mixture in a phytochamber (8 hr light/16 hr dark at 20° C and 18° C, respectively; 60 % humidity). See the supplemental methods for detailed information.

### Microscopic studies

Please refer to the supplementary experimental procedures for histology of anther, carpel, and spike development, as well as different microscopic methods like light, scanning electron, and fluorescence.

### Nucleic acid analysis

In the Supplemental methods, one can find methods for genomic DNA extraction, Southern hybridization, RNA extraction, and qRT-PCR.

### Nucleotide diversity (π) calculation

The whole-genome resequencing (WGS) data and SNP matrix for 200 diverse barley genotypes were downloaded from Jayakodi et al., 2020. The sequencing reads were aligned to the reference cv. Morex, as described (Jayakodi et al., 2020). The effectively covered areas of the barley genome were identified by the regions covered by at least two reads in ≥80% of the WGS accessions. The nucleotide diversity (π) was calculated on a 10 kb window with a step size of 2 kb with a custom script. Only the windows with ≥2 kb effectively covered region were considered. Please refer to the Supplemental methods for further nucleotide diversity analyses, including TILLING and resequencing of *HvHOX1* and *HvHOX2* in various genotypes and species.

### Microarray probe preparation and data analysis

The microarray probe preparation, hybridization, and data analysis were done as previously reported (Thirulogachandar et al., 2017). An elaborate method of the data preparation and co-expression network construction is given in the supplemental methods

### Data analysis

The qRT data were analyzed using the Prism software, version 8.4.2 (GraphPad Software, LLC). Mean value comparison of different traits was made with the multiple Student’s t-tests, paired Student’s t-test (parametric), and a one-way ANOVA with Tukey’s multiple comparison test (alpha=5%).

### Gene ontology enrichment analysis

The Gene Ontology (GO) enrichment analysis of differentially expressed genes and gene modules was done using the agriGO platform (v2) (Tian et al., 2017). The selected genes’ Arabidopsis IDs were queried against the Arabidopsis genome locus (TAIR9) reference set with the Fisher statistical test, Hochberg (FDR) multi-test adjustment method, and a significance level 0.05. The Plant GO slim “GO type” has been selected with a minimum number of entries. For final interpretation, the GO enrichment of biological processes was used.

### Transgenic and targeted mutagenesis

*In silico* identification of genes and promoters used for generating the transgenic plants used in this study are given in the Supplemental methods. Also, the methods of cloning various constructs, guide RNA design and preparation Cas9-triggered mutagenesis, as well as plant transformation are shown in the supplemental methods.

### Analysis of proteins

The preparation details of constructs used for the transactivation assay, electrophoretic mobility shift assay (EMSA), Western blot, and bimolecular fluorescence complementation assay are given in the supplemental methods.

## Author Contributions

V.T., N.S., T.S. G.G., and T.K. conceptualized the study. The study was supervised by N.S., T.S., and M.K. Microscopic analyses were done by V.T., and T.R. Transcriptome data were generated by V.T., and G.G., which was analyzed by V.T., and S.K. Transgenics and targeted gene-specific mutants were generated by G.H. and J.K. and analyzed by V.T. RGEN mutants were molecularly characterized by J.R. Resequencing of genes, and TILLING analysis were performed by R.K., T.S., S.S., T.K., and M.J. Constructs for protein characterization were prepared by G.G., V.T., P.S.R., and C.S. Transactivation and BiFC experiments were conducted by L.E-L., G.G., and J.L., and M.K. performed DNA binding study (EMSA). All the data were compiled, interpreted and drafted by V.T. The manuscript was reviewed by all the authors.

## Acknowledgments

We express our sincere gratitude to Jana Lorenz, Mandy Püffeld, Gabi Einert, Corinna Trautewig, and Angelika Püschel for their excellent technical support. We would like to thank Karin Lipfert, Heike Müller, Gudrun Schütze and Andreas Bähring for photography and graphical works. We are grateful to Sabine Sommerfeld und Sibylle Freist for technical assistance in barley transformation. We also extend our thanks to Dr. Nils Stein for providing access to TILLING screening platform and Stefan Ortleb for creating the spike movie. Work in the N.S. laboratory has been supported by a grant from the Ministry of Education Saxony-Anhalt (IZN), while work in the T.S. laboratory has been supported by the HEISENBERG Program of the German Research Foundation (DFG), grant nos. SCHN 768/8-1 and SCHN 768/15-1 and the IPK core budget.

## Accession number

*HvVHOX1* (*VRS1*) (Version: AB259782.1, GI: 119943316)

*HvHOX2* (Version: AB490233.1, GI: 266265607)

## Supplemental data

Supplemental Figure S1: Comparison of HvHOX1 and HvHOX2 nucleotide diversity in 200 domesticated barleys

Supplemental Figure S2: Pairwise alignment of HvHOX1 and HvHOX2 proteins

Supplemental Figure S3: Western blot for HvHOX1 and HvHOX2 proteins

Supplemental Figure S4: EMSA competition assay of in vitro translated HvHOX1 and HvHOX2

Supplemental Figure S5. Two-rowed spikes have delayed lateral spikelet development compared to its central spikelet and six-rowed lateral spikelets

Supplemental Figure S6. Transverse sections of anthers from central and lateral spikelets of Bowman and BW-NIL(vrs1.a)

Supplemental Figure S7: Transverse sections of carpels from central and lateral spikelets of Bowman and BW-NIL(vrs1.a)

Supplemental Figure S8: Comparison of HvHOX1 and HvHOX2 expression pattern during early spike development

Supplemental Figure S9. Hierarchial clustering of HvHOX1 and HvHOX2 shared and HvHOX1 unique modules

Supplemental Figure S10: Sequence alignment of the wild-type and mutant proteins of HvHOX1 and HvHOX2 resulted from the RGEN study

Supplemental Figure S11: The HvHOX2 gene might be a team player in barley spikelet development

Supplemental Figure S12: Multiple sequence alignment of the orthologous HvHOX2 proteins and HvHOX2 from a TILLING mutant 11869

Supplemental Figure S13. Gene ontology of Differentially expressed genes in W2.5, W3.0 and W4.5 in Bowman and BW-NIL(vrs1.a).

Supplemental Figure S14. Gene ontology of differentially expressed genes in W7.5 and W8.5 lateral spikelets of Bowman.

Supplemental Figure S15. Gene ontology of differentially expressed genes in W8.0 lateral spikelets of Transgenic plant E189 vs control plant E511.

Supplemental Methods. Additional methods and analyses used in this study.

Supplemental Table S1. HvHOX2 SNP haplotypes identified in 83 diverse spring barley collection

Supplemental Table S2. List of Hordeum species used in this study.

Supplemental Table S3. Nucleotide diversity of *HvHOX1* and *HvHOX2* in Hordeum species.

Supplemental Table S4. List of genes coexpressed with HvHOX1 and HvHOX2 genes during spike development in cv. Bowman

Supplemental Table S5. Primers used in this study.

